# Brain Structural Correlates of an Impending Initial Major Depressive Episode

**DOI:** 10.1101/2024.07.19.604266

**Authors:** Anna Kraus, Katharina Dohm, Tiana Borgers, Janik Goltermann, Dominik Grotegerd, Alexandra Winter, Katharina Thiel, Kira Flinkenflügel, Navid Schürmeyer, Tim Hahn, Simon Langer, Tilo Kircher, Igor Nenadić, Benjamin Straube, Hamidreza Jamalabadi, Nina Alexander, Andreas Jansen, Frederike Stein, Katharina Brosch, Paula Usemann, Lea Teutenberg, Florian Thomas-Odenthal, Susanne Meinert, Udo Dannlowski

## Abstract

**Background:** Neuroimaging research has yet to elucidate, whether reported gray matter volume (GMV) alterations in major depressive disorder (MDD) exist already before the onset of the first episode. Recruitment of presently healthy individuals with a known future transition to MDD (converters) is extremely challenging but crucial to gain insights into neurobiological vulnerability. Hence, we compared converters to patients with MDD and sustained healthy controls (HC) to distinguish pre-existing neurobiological markers from those emerging later in the course of depression.

**Methods:** Combining two clinical cohorts (*n*=1709), voxel-wise GMV of *n*=45 converters, *n*=748 patients with MDD, and *n*=916 HC were analyzed in regions-of-interest approaches. By contrasting the subgroups and considering both remission state and reported recurrence at a 2-year clinical follow-up, we stepwise disentangled effects of 1) vulnerability, 2) the acute depressive state, and 3) an initial vs. a recurrent episode.

**Results:** Analyses revealed higher amygdala GMV in converters relative to HC (*p*_TFCE-FWE_=.037, *d*=0.447) and patients (*p*_TFCE-FWE_=.005, *d*=0.508), remaining significant when compared to remitted patients with imminent recurrence. Lower GMV in the dorsolateral prefrontal cortex (*p*_TFCE-FWE_<.001, *d*=0.188) and insula (*p*_TFCE-FWE_=.010, *d*=0.186) emerged in patients relative to HC but not to converters, driven by patients with acute MDD.

**Conclusion:** By examining one of the largest available converter samples in psychiatric neuroimaging, this study allowed a first determination of neural markers for an impending initial depressive episode. Our findings suggest a temporary vulnerability, which in combination with other common risk factors might facilitate prediction and in turn improve prevention of depression.

## Introduction

Despite considerable effort, neuroimaging research has yet to unravel whether brain structural alterations in major depressive disorder (MDD) exist already before the onset of the initial depressive episode, indicating a neurobiological vulnerability. Meta-analyses of cross-sectional neuroimaging studies reported morphological changes in patients with MDD compared to healthy controls (HC; 5–7). Reduced gray matter volumes (GMV) in the dorsolateral prefrontal cortex (DLPFC), the insula, the rostral anterior cingulate cortex (rACC), and temporal gyrus are the most prominent findings in terms of cortical structural abnormalities in patients with MDD (2–5). Subcortically, the hippocampus and amygdala have been most extensively investigated with more heterogeneous findings (1,5,6). For the amygdala, reported alterations in GMV vary across studies and may depend on disease stage, psychopharmacological intake, and familiar risk for MDD (7–9). Due to the study designs and samples examined, these studies provide very limited insight into the temporal occurrence and persistence of GMV changes in MDD: are these alterations pre-existing and represent a neural vulnerability for the future onset of the disorder or do they emerge later as a correlate of MDD developing in the course of the disorder? And further, are changes in GMV of persistent nature or do they follow the dynamic course of the disorder, e.g., emerging only during acute depressive episodes and disappearing in times of remission (13)?

Consequently, we lack a consistent neurobiological theory to explain the complex interplay between depression and brain structure. Available studies showed less GMV in MDD in cortical and limbic structures related to the number of recurrent episodes, duration of illness (6,11–14) or longitudinally volume decline as a function of relapse (15–17) or rehospitalization (18). Supporting the assumption of a rather dynamic correlate, more amygdala GMV has been reported in association with a current first depressive episode (19,20) while longitudinal studies found support for state-like volume decrease in both cortical and subcortical regions driven by current depressive symptoms (15,21,22). There is a crucial gap in our understanding of how GMV changes may indicate a marker for the future onset of MDD in presently healthy individuals. To shed light on neurobiological vulnerability, we need to examine individuals with a known future transition to MDD (converters) as this is indispensable to determine pre-existing changes prior to the onset of MDD (i.e., the onset of a first episode). We face a staggering lack of studies in converter samples, while the little available research was conducted mainly on children and adolescents. These studies found reduced GMV in frontal and temporal areas, insula, and rACC to be associated with the future onset of MDD or rise in depressive symptoms (23,24). Two studies reported enhanced amygdala volumes (25,26) in association with impending transition to MDD. However, besides low quality diagnostic measurement (via questionnaire-based self-reports, rather than clinical ratings) and the lack of control for confounding effects (e.g., due to pubertal changes or family risk), these studies are drastically underpowered (e.g., average converter sample size of n=22.55; (24). Small sample sizes are hardly surprising as recruitment of converters is exceptionally challenging: Considering the annual incidence of MDD of 4.3% (27), this requires the inclusion of large numbers of HC undergoing neuroimaging, alongside extensive long-term clinical follow-up assessments, to post-hoc identify an adequately sized sample of converters. Moreover, to the best of our knowledge, no available study has conducted a comparison of structural imaging data between converters and already affected patients with MDD in different remission states.

We faced this challenge by analyzing voxel-wise GMV of 45 initially healthy individuals with a known future transition to MDD (converters), healthy individuals who do not develop a MDD within the next two years and a large heterogeneous sample of patients with MDD. Based on literature reporting that the DLPFC, insula, rACC as well as the hippocampus and amygdala seem to be particularly affected in MDD, a region of interest (ROI) approach was utilized with additional exploratory whole-brain analyses. By contrasting converters to patients with MDD and HC, we aimed at disentangling GMV alterations in MDD in markers of vulnerability for an impending onset of an initial depressive episode and neural correlates of MDD that emerge later in the course of the disorder (analysis 1). To further investigate the impact of the acute depressive state (i.e., severe depressive symptoms during an acute depressive episode) on GMV, we subsequently only included patients in remission (analysis 2). Lastly, we compared converters with patients in remission and known recurrence within the follow-up period to examine neural underpinnings of an *initial* depressive episode vs. a recurrent episode (analysis 3).

## Methods and Materials

### Sample

The present study combined two independent cohorts: the Marburg-Münster Affective Disorders Cohort Study (MACS) and the Münster Neuroimaging Cohort (MNC). Both ongoing studies comprise converters, patients with MDD and HC without transition to any mental disorder after follow-ups. Participants were aged between 18 and 65 years at baseline. For all participants, a magnetic resonance imaging (MRI) assessment and a consecutive clinical follow-up after approximately two years were considered. Individuals in the converter sample and HC showed absence of any psychiatric diagnosis according to DSM-IV-TR at MRI assessment but converters fulfilled the criteria for acute or lifetime MDD at the 2-year clinical follow-up. For details on recruitment and exclusion criteria see Supplementary Methods.

From the MACS, *n*=1279 participants were analyzed, consisting of *n*=30 converters, *n*=590 patients with MDD and *n*=659 HC. MRI data collection for this study took place between September 2014 and June 2019. Patients diagnosed with MDD in the MACS cohort showed varying levels of symptom severity and underwent a range of treatments (inpatient, outpatient, or none). The MACS was conducted at two scanning-sites: University of Muenster and University of Marburg (see 31 for the general study description and 32 for MRI quality assurance protocol).

From the MNC, *n*=430 participants were included in this study, consisting of *n*=15 converters, *n*=158 patients with MDD and *n*=257 HC. MRI data was acquired between October 2009 and December 2020. Patients were suffering from a moderate or severe depressive episode and underwent inpatient treatment at MRI assessment.

The three groups did not differ regarding age or sex in the total sample (Table 1). Converters showed a significantly higher ratio of familial risk for MDD than HC but no differences in depressive symptoms. Compared to patients with MDD, converters experienced less depressive symptoms. Patients with MDD in the MNC reported a significantly higher number of lifetime in-patient treatments and higher levels of psychotropic medication intake compared to patients in the MACS (Table 2).

**Table 1.** Sample characteristics of the total sample, the remission sample and the recurrence sample by group.

| Characteristic | Total sample (*n*=1709) |  |  |  |  |  |  | Remission sample (*n*=1310)<sup>a</sup> |  |  |  | Recurrence sample (*n*=1109)<sup>b</sup> |  |  |  |
| --- | --- | --- | --- | --- | --- | --- | --- | --- | --- | --- | --- | --- | --- | --- | --- |
|  | CON<br>(*n*=45) | HC<br>(*n*=916) | MDD<br>(*n*=748) |  | CON<br>vs.<br>HC | CON<br>vs.<br>MDD | HC<br>vs.<br>MDD | MDD<br>(*n*=349) |  | CON<br>vs.<br>MDD | HC<br>vs.<br>MDD | MDD<br>(*n*=148) |  | CON<br>vs.<br>MDD | HC<br>vs.<br>MDD |
|  | *M* (*SD*) | *M* (*SD*) | *M* (*SD*) | *p*<sup>1</sup> | *p*<sup>2</sup> | *p*<sup>2</sup> | *p*<sup>2</sup> | *M* (*SD*) | *p*<sup>1</sup> | *p*<sup>2</sup> | *p*<sup>2</sup> | *M* (*SD*) | *p*<sup>1</sup> | *p*<sup>2</sup> | *p*<sup>2</sup> |
| Age | 34.02<br>(11.46) | 36.45<br>(13.04) | 37.44<br>(13.29) | .108 | .174 | .059 | .127 | 37.61<br>(13.23) | .141 | .083 | .160 | 35.73<br>(13.44) | .413 | .440 | .536 |
| Gender (m/f)<sup>3</sup> | 18/27 | 372/544 | 273/475 | .227 | .935 | .636 | .087 | 112/237 | **.020** | .288 | **.005** | 44/104 | **.042** | .196 | **.012** |
| Cohort (MACS/<br>MNC)<sup>3</sup> | 30/15 | 659/257 | 590/158 | .002 | .443 | .054 | **.001** | 344/5 | **<.001** | **<.001** | **<.001** | 148/0 | **<.001** | **<.001** | **<.001** |
| HDRS Score | 1.71<br>(2.50) | 1.16<br>(1.73) | 10.29<br>(7.41) | **<.001** | .151 | **<.001** | **<.001** | 4.99<br>(4.61) | **<.001** | **<.001** | **<.001** | 6.04<br>(4.72) | **<.001** | **<.001** | **<.001** |
| Familiar risk for<br>MDD (no<br>risk/risk)<sup>3</sup> | 31/14 | 786/124 | 553/189 | **<.001** | **.001** | .401 | **<.001** | 234/114 | **<.001** | .824 | **<.001** | 99/49 | **<.001** | .802 | **<.001** |

CON=converters, HC=healthy controls, MDD=major depressive disorder, MACS=Marburg Münster Affective Disorders Cohort Study, MNC=Münster Neuroimaging Cohort, HDRS=Hamilton Depression Rating Scale. Significant *p*-values (*p*<.05) are highlighted in bold.
<sup>a</sup> characteristics of CON and HC see total sample. Statistic parameters of CON vs. HC comparison equals the total sample. Includes patients with remitted MDD only.
<sup>b</sup> characteristics of CON and HC see total sample. Includes patients with remitted MDD with recurrence within 2-year follow-up interval only.
<sup>1</sup> MDD vs. CON vs. HC using one-way ANOVA except where noted.
<sup>2</sup> unpaired two-tailed *t*-test except where noted.
<sup>3</sup> X<sup>2</sup>-test.

**Table 2.** Clinical characteristics of patients with MDD and converters for the total sample, the remission sample and the recurrence sample by study cohort.

| Characteristic | Patients with Major Depressive Disorder |  |  |  |  |  |  |  |  | Converters |  |  |
| --- | --- | --- | --- | --- | --- | --- | --- | --- | --- | --- | --- | --- |
|  | Total sample |  |  | Remission sample<sup>a</sup> |  |  | Recurrence sample<sup>b</sup> |  |  | MACS<br>(*n*=30) | MNC<br>(*n*=15) | *p*<sup>1</sup> |
|  | MACS<br>(*n*=590) | MNC<br>(*n*=158) | *p*<sup>1</sup> | MACS<br>(*n*=344) | MNC<br>(*n*=5) | *p*<sup>1</sup> | MACS<br>(*n*=148) | MNC<br>(*n*=0) | *p*<sup>1</sup> |  |  |  |
|  | *M*<br>(*SD*) | *M*<br>(*SD*) |  | *M*<br>(*SD*) | *M*<br>(*SD*) |  | *M*<br>(*SD*) | *M*<br>(*SD*) |  | *M*<br>(*SD*) |  |  |
| Time since onset (month) | 131.92<br>(121.50) | 128.09<br>(124.98) | .739 | 137.31<br>(120.80) | NA | NA | 143.19<br>(130.40) | NA | NA | NA | NA | NA |
| Number of depressive episodes lifetime | 3.81<br>(6.58) | 4.60<br>(5.79) | .173 | 3.26<br>(5.29) | 1.80<br>(4.03) | .540 | 4.01<br>(6.74) | NA | NA | NA | NA | NA |
| Cumulative duration of depression (month) | 43.61<br>(61.92) | 34.97<br>(48.69) | .070 | 38.16<br>(55.56) | 24.00<br>(53.67) | .572 | 46.86<br>(71.91) | NA | NA | NA | NA | NA |
| Number of in-patient treatments lifetime | 1.48<br>(1.93) | 2.13<br>(1.72) | **<.001** | 1.19<br>(1.67) | .40<br>(0.89) | .294 | 1.26<br>(1.50) | NA | NA | NA | NA | NA |
| Cumulative duration of in-patient treatment (weeks) | 11.79<br>(18.10) | 11.95<br>(16.03) | .918 | 10.09<br>(17.65) | 2.40<br>(5.37) | .332 | 9.04<br>(10.71) | NA | NA | NA | NA | NA |
| Comorbidities (no/yes)<sup>2</sup> | 336/253 | 78/80 | .085 | 209/135 | 4/1 | .381 | 80/68 | NA | NA | NA | NA | NA |
| Medication Load Index | 1.32<br>(1.49) | 2.46<br>(1.70) | **<.001** | .95<br>(1.18) | .40<br>(0.89) | .295 | 1.05<br>(1.11) | NA | NA | NA | NA | NA |
| Time until onset (month) | NA | NA | NA | NA | NA | NA | NA | NA | NA | 12.29<br>(7.29) | 8.36<br>(7.22) | .116 |

MACS=Marburg Münster Affective Disorders Cohort Study, MNC=Münster Neuroimaging Cohort. Significant p-values ( $p < .05$ ) are highlighted in bold.
<sup>a</sup> includes partly or fully remitted patients only.
<sup>b</sup> includes partly or fully remitted patients only who experienced a relapse within follow-up interval.
<sup>1</sup> MACS vs. MNC using unpaired two-tailed t-test except where noted.
<sup>2</sup> X<sup>2</sup>-test

The MACS was approved by the Ethics Committees of the Medical Faculties, University of Marburg (07/2014) and University of Muenster (2014-422-b-S). The MNC was approved by the ethics committee of the Medical Faculty of University of Muenster (2007-307-f-S). All participants provided written informed consent and received financial compensation.

### Materials and Procedure

All participants underwent structural MRI. The Structural Clinical Interview for DSM-IV-TR (SCID-IV) (30) was conducted by trained personnel to verify presence or absence of psychiatric diagnosis at MRI assessment and clinical follow-up. At clinical follow-up, a life-chart method was used to accurately assess individual disease courses (31).

Current depression severity was assessed using the Hamilton Depression Rating Scale (HDRS; 35). To determine familial risk, participants were asked whether they had a first-degree relative who had been diagnosed with an affective disorder and received treatment for this. To account for psychotropic medication intake, a medication load index (MedIndex) was calculated as utilized in previous studies (18,37,38, see Supplementary Methods).

### Structural image acquisition and preprocessing

T1-weighted high-resolution anatomical images were acquired at 3T MRI scanners through three-dimensional fast gradient echo sequences (MACS: MPRAGE, MNC: turbo field echo). Details on acquisition parameters can be found in Supplementary Methods for both cohorts separately. Within the MRI assessment period in the MACS cohort, the body-coil at the Marburg site was exchanged, which resulted in two dummy-coded variables considered by the applied harmonization algorithm. Further information on quality assurance protocol and scanner harmonization can be found elsewhere (29).

All T1-weighted images were preprocessed together using an identical processing and quality checking pipeline, employing the default parameter settings of the CAT12 toolbox (www.neuro.uni-jena.de/cat, v1720) implemented in the Statistical Parametric Mapping software (SPM12, version 7771, Institute of Neurology, London, UK). Briefly, processing steps included bias-correction, tissue segmentation, and spatially normalization to MNI-space using linear (12-parameter affine) and non-linear transformations, within a unified model including high-dimensional geodesic shooting normalization (35). Gray matter images were smoothed with an 8 mm full width at half-maximum Gaussian kernel and underwent outlier detection using CAT12’s check homogeneity function and visual inspections. To adjust for scanner and site effects, data was harmonized using the batch correction tool ComBat (39, see Supplementary Methods).

### Statistical analyses

IBM SPSS Statistics 28 (SPSS Inc., Chicago, IL, USA) was utilized to analyze the descriptive and clinical data. Plots were created in R (https://www.R-project.org/). All second-level analyses of voxel-wise MRI data were performed with SPM12. We selected the rostral middle frontal gyrus, the insula, the rACC (subgenual and pregenual), the amygdala, and the hippocampus as ROIs and created separate bilateral masks according to the automated anatomical labelling atlas 3 (AAL3; 44). The AAL3 is integrated into the Wake Forest University pickatlas (38) in SPM12. Following earlier studies (16,17), the middle frontal gyrus was chosen as representative of the DLPFC, since it encompasses Brodmann’s area 46 (39). All statistical analyses were performed separately for each ROI. Bonferroni correction was applied to account for multiple statistical tests across *n*=5 ROIs (implying a threshold of *p*=.01). ROI analyses were conducted with a threshold-free cluster enhancement (TFCE) implemented in the TFCE-toolbox (http://dbm.neuro.uni-jena.de/tfce, Version222) with 5000 permutations per test and an absolute threshold masking of 0.1. The statistical threshold was set to a conservative familywise-error (FWE) correction of *p*<0.05 on the voxel-level. Exploratory whole-brain analyses were performed at a statistical threshold of *p*<0.001, uncorrected, and a cluster threshold of *k*>200 voxels.

#### Analysis 1: vulnerability

To investigate whether GMV changes represent a vulnerability, we performed full-factorial ANCOVAs on gray matter segments with group (converters vs. MDD vs. HC) as between-subjects factor and age, sex, and total intracranial volume (TIV) as covariates of no interest for each ROI separately and post-hoc two-sided *t*-tests in case of significance. Any GMV alterations in converters compared to HC would indicate markers of the impending onset of MDD, representing either temporary vulnerability if emerging in converters only or rather stable pre-existing alterations if both converters and patients with MDD deviate from HC.

#### Analysis 2: effects of acute depressive state

To distinguish previously identified effects in MDD from effects of an acute depressive state (e.g., severe depressive symptoms during an acute depressive episode), significant *t*-tests from analysis 1 were subsequently recalculated in a sample, which only comprised patients with MDD in partial or full remission (remission sample). Consequently, any remaining deviations in patients with remitted MDD from converters and HC would indicate rather persisting neural correlates of MDD.

#### Analysis 3: first episode vs. recurrence

To disentangle whether effects are present only before the first-time depressive episode or also prior a recurrent episode, significant *t*-tests from analysis 2 were further recalculated in a sample that included patients with currently remitted MDD, who furthermore experienced recurrence within the follow-up interval (recurrence sample). A recurrent episode was assessed the identical way as the converters’ first episode within the follow-up interval.

#### Additional Analyses

To address potential impact of large size differences of the subgroups on significant results, we additionally repeated the procedure described in analysis 3 in a smaller sample matched for age, sex, and scanner variables in a last step (see Supplementary Methods). Further sanity checks (e.g., effects of confounding variables, outliers, time until onset of initial depressive episode in converter) can be found in the Supplementary Methods.

## Results

### Analysis 1: vulnerability

ROI analyses in the DLPFC, insula and amygdala revealed significant main effects of group (all *p*_TFCE-FWE_<.048, partial η²>.007) but not in the rACC (*p*_TFCE-FWE=_.184) and hippocampus (*p*_TFCE-FWE=_.142). While the significant effects were mainly driven by lower GMV in patients with MDD compared to HC (all *p*_TFCE-FWE_<.010, *d*>.186), in the DLPFC and insula (Figure S1 in Supplementary Results), we found significantly higher amygdala volumes in converters compared to both HC (*t*(1703)=2.93, *p*_TFCE-FWE_ =.037, Cohen’s *d*=0.447; Figure 1A) and patients with MDD (*t*(1703)=3.31, *p*_TFCE-FWE_ =.005, Cohen’s *d*=0.508, Figure 1B). In the DLPFC and insula, no differences emerged between patients with MDD compared to converters or in converters compared to HC in post-hoc *t*-tests. For details, see Supplementary Results.

**Figure 1.**
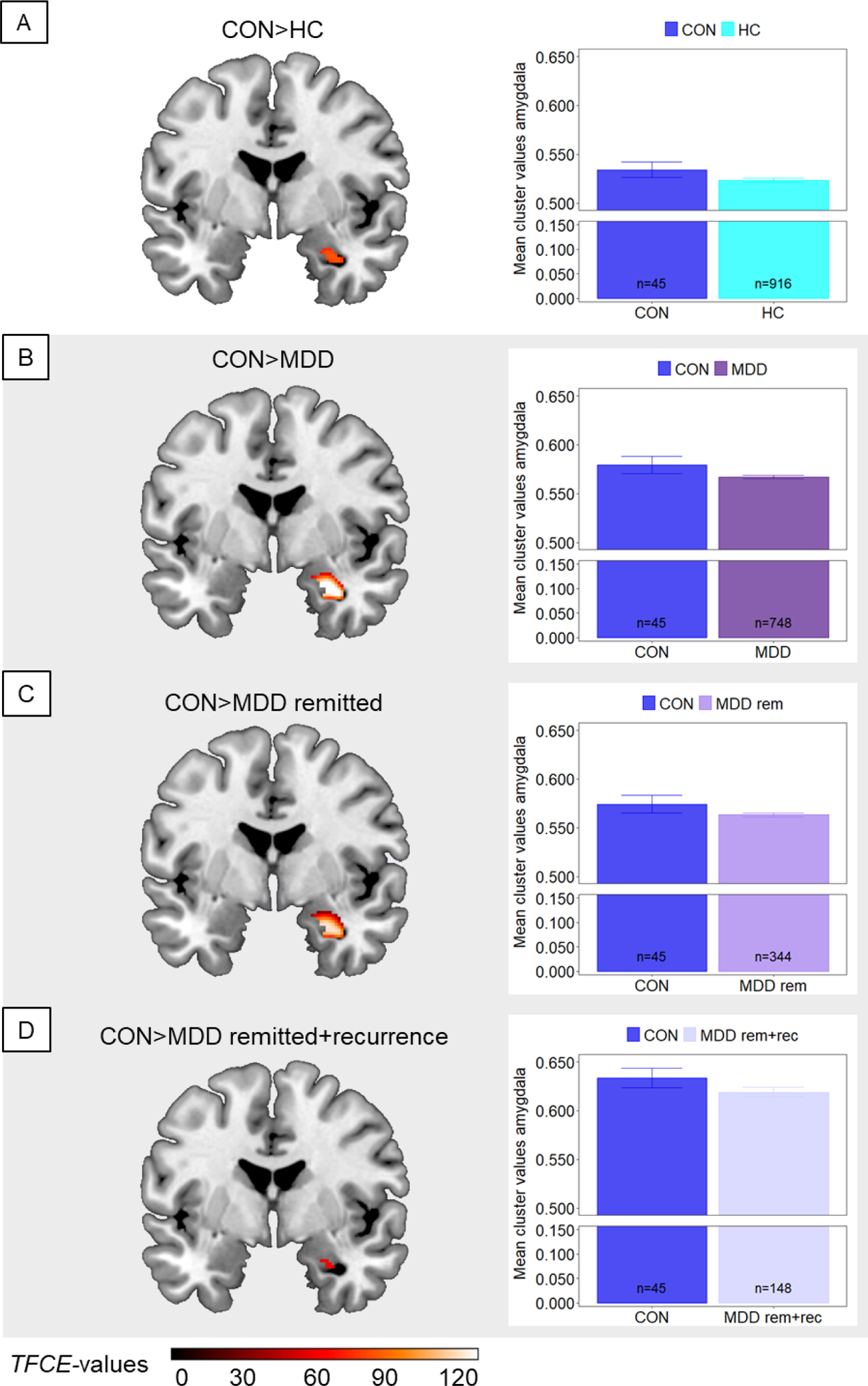
Larger gray matter volumes in the amygdala in converters compared to HC and patients with MDD in different remission states. Comparisons between converters (CON) and subgroups are shown. Clusters are thresholded at *p*_TFCE-FWE_<.05 and bar plots represent respective extracted Eigenvariates from the depicted cluster. Error bars indicate 1 SEM. *T*-contrasts are shown for the comparison of converters with A) healthy controls (HC), B) patients with major depressive disorder (MDD) in heterogeneous remission states, C) patients with currently remitted MDD (MDD rem), and D) patients with currently remitted MDD with recurrence within a 2-year clinical follow-up period (MDD rem+rec). Shaded subplots indicate comparisons of converters with MDD subgroups.

### Analysis 2: effects of acute depressive state

In ROI analyses in the remission sample, the *t*-tests did not reveal lower GMV in the DLPFC (*p*_TFCE-FWE=_.114) and insula (*p*_TFCE-FWE=_.175) in patients with remitted MDD compared to HC. In the amygdala, however, converters showed more GMV compared to patients with remitted MDD (*t*(1299)=3.17, *p*_TFCE-FWE_ =.002, Cohen’s *d*=0.502, Figure 1C) and compared to HC (*t*(1299)=2.86, *p*_TFCE-FWE_ =.014, Cohen’s *d*=0.437). For details, see Supplementary Results.

### Analysis 3: first episode vs. recurrence

Since in analysis 2, we found significant effects only in the amygdala, we repeated the ROI analysis in the amygdala in the recurrence sample. Again, the *t*-tests revealed larger volumes in the amygdala in converters compared to patients with remitted MDD and known recurrence within the follow-up interval (*t*(1103)=2.67, *p*_TFCE-FWE=_.046, Cohen’s *d*=0.455; Figure 1D) as well as compared to HC (*t*(1103)=2.83, *p*_TFCE-FWE=_.024, Cohen’s *d*=0.432). There were no significantly lower amygdala volumes in patients compared to HC (*p*_TFCE-FWE_=.629).

### Additional analyses

Findings from exploratory whole-brain analyses in the three samples concordant with reported effects from the ROI analyses in the DLPFC and insula in clusters comprising e.g., temporal and frontal regions, thalamus and insula (see Supplementary Results).

For detailed results on sanity checks investigating effects of large sample size differences, confounding variables (e.g., psychiatric medication or familiar risk for MDD), outliers, or the time until onset of the initial depressive episode in the converter subsample see Supplementary Results. Briefly, the amygdala effect remained significant after Bonferroni correction for multiple testing, independent of added covariates, and in the smaller sample matched for age, sex, and scanner variables. After controlling for familiar risk for MDD, group differences described in analysis 1 remained significant. When adding medication as covariate, remaining significant effects were comparable to results described in analysis 2.

## Discussion

This study investigated cross-sectional differences in GMV of initially healthy individuals with a known future transition to MDD (converters), patients with diagnosed MDD in different remission states, and HC with no previous and also no future depressed episodes within an identical follow-up period. Aim of the study was to disentangle pre-existing markers of neurobiological vulnerability for the future onset of a depressive episode and correlates of MDD emerging in already affected individuals, especially focusing on the impact of the acute depressive state. Two main findings emerged from analyses: firstly, converters showed more GMV in the amygdala compared to patients with MDD and HC. Since this effect remained significant when contrasting converters and remitted patients with known recurrence within the follow-up period, the amygdala enlargement could be a temporary marker specific for the future onset of MDD (i.e., the onset of a first episode). Secondly, less volumes in the DLPFC and insula in patients with MDD most likely do not represent vulnerability for depression and are at least partially driven by the acute depressive state.

The observed morphometric changes in the amygdala, insula and prefrontal cortex correspond to neurobiological models of MDD assuming a dysfunction in the frontolimbic brain circuitry (40). Putatively, this manifests through an inhibition of top-down mechanisms with reduced activity of the DLPFC, which leads to impaired cognitive control over emotional reactions. At the same time, increased bottom-up processing through enhanced amygdala and insula activity to negative stimuli contributes to difficulties in emotion processing, which may exacerbate depressive symptoms (40–43).

We found more GMV in the amygdala in converters relative to both patients with MDD and HC. The converter sample comprises currently healthy individuals who do not yet show elevated levels of depressive symptoms compared to HC but are known to experience their first-time depressive episode within the next two years. Thus, the amygdala enlargement neither reflects a state effect associated to acute depressive symptoms nor a persistent neural correlate of depression. It might rather represent a temporary vulnerable state that marks the onset of an impending first depressive episode. Findings from longitudinal research are mostly inconsistent regarding amygdala GMV in association with onset (24) as well as recurrence (21,44) of depression. Most of these studies, along with our results, converge in suggesting that amygdala volume may track the dynamic course of the disorder, which in turn would reflect the amygdala’s key role in generation and stability of affective states (7,43). To date, the precise trajectory of these alterations and their associations with further risk factors (e.g., environmental factors) remains unclear. Observations of more amygdala GMV in converters relative to HC align with Nickson et al. (2016). Although their study lacked a direct comparison with a clinical MDD sample, they reported a decline in amygdala volume over time in individuals who developed MDD, a pattern that corresponds with our findings of reduced amygdala volumes within patients. Research in patients with acute first episode depression frequently reported enlarged amygdala volumes (19,20), with one study also noting this when contrasting with recurrent MDD (45). Taken together with our findings from the comparison of converters and remitted patients with known recurrence within follow-up, the amygdala enlargement may be a neural marker of the impending *onset* of MDD, rather than a recurrent episode. Potentially, this could be associated with enhanced metabolism in the amygdala and persistent hyperactivity to negative stimuli in MDD (46,47). As the disease progresses, stress-related neurotoxic processes (48–51) during recurrent depressive episodes might lead to a subsequent decline in GMV. Thus, alterations related to the disease stage could moreover serve as an explanation for the very heterogeneous findings on amygdala volume in the existing literature.

Our findings on lower GMV in the DLPFC and insula ROIs in patients with MDD, but also in frontal and temporal regions that emerged in whole-brain analyses, do not seem to indicate vulnerability for MDD, since we did not observe such alterations in the comparison of converters and HC. Furthermore, when only including patients with currently remitted MDD in the analyses, effects were no longer significant. Our results suggest that lower volumes in the DLPFC and insula may reflect the acute depressive state to some extent and partially normalize with remission rather than being a persistent neural correlate of MDD. In our recent work on replicability and generalizability of gray matter alterations in MDD we found a rather similar pattern of GMV reductions across stratified subsamples of patients with MDD (e.g., acute/remitted, medicated/non-medicated) implying a general MDD diagnosis effect (5). Since our remission sample was restricted to a lower sample size of remitted patients (*n*=349), our results could be driven by insufficient statistical power considering small effect sizes in neuroimaging modalities (52). Nevertheless, replicability was highest when contrasting HC with acutely depressed patients (5), which could be related to effects of the acute depressive state to some extent. Previous cross-sectional (13,53,54) and longitudinal research (18,55) on volumes of DLPFC and insula suggested neural correlates of MDD might actually represent a persistent, even scar-like effect, as stronger GMV decline was associated with more severe disease courses (e.g., duration of disease or number of episodes). However, evidence on such scar-like effects is limited due to cross-sectional study designs and in general the inadequately powered sample sizes of patient (sub)groups coming with a risk for inflated effects (56). In line with our results, longitudinal studies found GMV reductions in patients to disappear with statistical control for influence of current depression severity and that GMV in times of remission either remained unchanged while it decreased in non-remitters (15,17) or even increased (16). Additional preliminary evidence on the impact of acute depressive state on the brain comes from recent studies in other neuroimaging modalities such as task-based fMRI (57) and structural connectome analysis (58), though there is little research on the direct link between modalities. Less severe depressive symptoms, e.g., in times of remission, might be associated with less subjective stress exposure, related to the inhibition of stress-mediated neurotoxic processes in depression (48–51) in structures of the frontolimbic circuitry. Following this, presumably increased stress levels in converters before they experience their first-time depressive episode might also explain the absence of volume differences in the DLPFC or insula between converters and patients with MDD as well as between converters and HC. Potentially, an incipient GMV decline in these reciprocally connected structures (59) may be present albeit too small to achieve statistical significance in the converter sample.

Regarding GMV of the hippocampus and rACC, we did not find any significant effects between the groups, which stands in contrast to earlier research (1–3). Nevertheless, our null-findings for the hippocampus are consistent with two recent studies both conducted in samples comprising over 4000 individuals (5,60).

The major strength of this study is the unprecedented large sample of individuals with a known future transition to MDD, which required the recruitment of a large number of HC undergoing neuroimaging with additional clinical follow-up assessments. Data of this valuable population is crucial to make assumptions about neuronal vulnerability for the onset of MDD. Contrasting converters and HC with patients in different states of remission also expanded our understanding of the impact of the acute depressive state on GMV. Another strength is that we analyzed the characteristics of the first-time depressive episode according to SCID-VI (30) as opposed to a recurrent episode. Moreover, we accounted for confounding effects of size differences of subsamples, psychopharmacological treatment and familiar risk for MDD.

A few limitations need to be addressed. Firstly, the subgroups strongly differed in size and considering our recent findings (52), the size of the converter sample is insufficient to detect small effects common in neuroimaging modalities. To account for size differences, we created a subsample matched for sex, age and scanner variables. Moreover, compared to previous imaging studies in converters (24,61), our sample (*n*=45) is to our knowledge among the largest so far. Secondly, the time until onset of MDD in the converters varied between one and 24 months, so no certain conclusions can be drawn about the exact beginning of the reported morphometric alterations. Perhaps stronger effects could be observed with a shorter time until onset. In this sample, however, the GMV of amygdala, insula and DLPFC was not associated with the months until onset. Thirdly, we combined two clinical cohorts using different scanners to increase sample size. Since in the MNC almost all patients were acutely depressed, the remission and the recurrence sample comprised mainly remitted patients from the MACS cohort whereas converters and HC were included from both cohorts. To address potential scanner and cohort effects in the remission sample, data was harmonized in the total sample using ComBat (39) prior excluding acute cases.

In conclusion, examining one of the largest available converter samples in neuroimaging research offered preliminary evidence on neural markers for the impending onset of an initial depressive episode and other correlates of MDD that emerge later in the course of the disorder and are probably driven by the acute depressive state. Our findings extend neurobiological models of the etiology of MDD (40–43) and suggest a temporary vulnerability, which in combination with other common risk factors for MDD might facilitate prediction of the onset of the disorder. In turn, this could improve both early detection and prevention programs for depression. Finally, encouragement is given for longitudinal studies focusing on such highly relevant converter samples prior to, during, and following the initial depressive episode to further elucidate GMV trajectories in the early stages of depression.

## Acknowledgements

This work is part of the German multicenter consortium “Neurobiology of Affective Disorders. A translational perspective on brain structure and function“, funded by the consortia grants from the German Research Foundation (Deutsche Forschungsgemeinschaft DFG; Forschungsgruppe/Research Unit FOR2107) FOR 2107 and SFB/TRR 393 (grant FOR2107 KI588/14-1, KI588/14-2,, KI588/15-1, KI588/17-1, KI588/20-1, KI588/22-1 to TK, DA1151/5-1, DA1151/5-2, DA1151/6-1, DA1151/9-1, DA1151/10-1, DA1151/11-1 to UD, STR1146/18-1 to BS, NE2254/1-2, NE2254/2-1, NE2254/3-1, NE2254/4-1, JA1890/7-1, JA1890/7-2 to AJ, HA7070/2-2 to TH; grant SFB-TRR393, Projects A01 and S03 to TH, A02 and Z to TK, A02 and S02 to UD; A04 to SM, A04 and C02 to IN, B01 and INF to AJ, B03 and RTG to BS, B03 and S03 to HJ, B05 and S02 to NA, INF to FS), the Interdisciplinary Center for Clinical Research (IZKF) of the medical faculty of Münster (grant Dan3/022/22 to UD) and the “Innovative Medizinische Forschung” (IMF) of the medical faculty Münster (grant ME122205 to SM; grant KO-121806 to KD).

The research unit FOR 2107 is divided into the following Working Packages (WP):

WP1, FOR2107/MACS cohort and brainimaging. Authors include: TK (speaker FOR2107), UD (co-speaker FOR2107), IN (principal investigator (PI); JG, AW, KT, KF, DG, FS, KB, PU, LT, FT-O, SM. Other non-author PIs and members include: Axel Krug (PI; KR 3822/5-1, KR 3822/7-2), Carsten Konrad (PI; KO 4291/3-1), Henrike Bröhl, Bruno Dietsche, Rozbeh Elahi, Jennifer Engelen, Ulrika Evermann, Sabine Fischer, Jessica Heinen, Svenja Klingel, Felicitas Meier, Tina Meller, Julia-Katharina Pfarr, Kai Ringwald, Torsten Sauder, Simon Schmitt, Annette Tittmar, Adrian Wroblewski, Dilara Yüksel (Dept. of Psychiatry, Marburg University). Mechthild Wallnig, Rita Werner (Core-Facility Brainimaging, Marburg University). Carmen Schade-Brittinger, Maik Hahmann (Coordinating Centre for Clinical Trials, Marburg). Michael Putzke (Psychiatric Hospital, Friedberg). Rolf Speier, Lutz Lenhard (Psychiatric Hospital, Haina). Birgit Köhnlein (Psychiatric Practice, Marburg). Peter Wulf, Jürgen Kleebach, Achim Becker (Psychiatric Hospital Hephata, Schwalmstadt-Treysa). Ruth Bär (Care facility Bischoff, Neukirchen). Matthias Müller, Michael Franz, Siegfried Scharmann, Anja Haag, Kristina Spenner, Ulrich Ohlenschläger (Psychiatric Hospital Vitos, Marburg). Matthias Müller, Michael Franz, Bernd Kundermann (Psychiatric Hospital Vitos, Gießen). Christian Bürger, Fanni Dzvonyar, Verena Enneking, Stella Fingas, Katharina Förster, Hannah Lemke, Nils Opel, Ronny Redlich, Jonathan Repple, Kordula Vorspohl, Bettina Walden, Lena Waltemate, Dario Zaremba (Dept. of Psychiatry, University of Münster). Harald Kugel, Jochen Bauer, Walter Heindel, Birgit Vahrenkamp (Dept. of Clinical Radiology, University of Münster). Gereon Heuft, Gudrun Schneider (Dept. of Psychosomatics and Psychotherapy, University of Münster). Thomas Reker (LWL-Hospital Münster). Gisela Bartling (IPP Münster). Ulrike Buhlmann (Dept. of Clinical Psychology, University of Münster).

WP2, animal phenotyping. Non-author PIs and members include: Markus Wöhr (PI; WO 1732/4-1, WO 1732/4-2), Rainer Schwarting (PI; SCHW 559/14-1, SCHW 559/14-2)Marco Bartz, Miriam Becker, Christine Blöcher, Annuska Berz, Moria Braun, Ingmar Conell, Debora dalla Vecchia, Darius Dietrich, Ezgi Esen, Sophia Estel, Jens Hensen, Ruhkshona Kayumova, Theresa Kisko, Rebekka Obermeier, Anika Pützer, Nivethini Sangarapillai, Özge Sungur, Clara Raithel, Tobias Redecker, Vanessa Sandermann, Finnja Schramm, Linda Tempel, Natalie Vermehren, Jakob Vörckel, Stephan Weingarten, Maria Willadsen, Cüneyt Yildiz (Faculty of Psychology, Marburg University).

WP3, miRNA: Non-author PI: Gerhard Schratt (SCHR 1136/3-1, 1136/3-2).

WP4, immunology, mitochondriae. Non-author PIs and members include: Judith Alferink (PI; AL 1145/5-2), Carsten Culmsee (PI; CU 43/9-1, CU 43/9-2), Holger Garn (PI; GA 545/5-1, GA 545/7-2), Jana Freff (Dept. of Psychiatry, University of Münster). Susanne Michels, Goutham Ganjam, Katharina Elsässer (Faculty of Pharmacy, Marburg University). Felix Ruben Picard, Nicole Löwer, Thomas Ruppersberg (Institute of Laboratory Medicine and Pathobiochemistry, Marburg University).

WP5, genetics. Non-author PIs and members include: Marcella Rietschel (PI; RI 908/11-1, RI 908/11-2), Markus Nöthen (PI; NO 246/10-1, NO 246/10-2), Stephanie Witt (PI; WI 3439/3-1, WI 3439/3-2), Helene Dukal, Christine Hohmeyer, Lennard Stütz, Viola Lahr, Fabian Streit, Josef Frank, Lea Sirignano (Dept. of Genetic Epidemiology, Central Institute of Mental Health, Medical Faculty Mannheim, Heidelberg University). Stefanie Heilmann-Heimbach, Stefan Herms, Per Hoffmann (Institute of Human Genetics, University of Bonn, School of Medicine & University Hospital Bonn). Andreas J. Forstner (Institute of Human Genetics, University of Bonn, School of Medicine & University Hospital Bonn; Centre for Human Genetics, Marburg University).

WP6, multi-method data analytics. Authors include: TH (PI; HA 7070/2-2). Non-author PIs and members include: Bertram Müller-Myhsok (PI; MU1315/8-2), Astrid Dempfle (PI; DE 1614/3-1, DE 1614/3-2), Anastasia Benedyk, Miriam Bopp, Roman Keßler, Maximilian Lückel, Verena Schuster, Christoph Vogelbacher (Dept. of Psychiatry, Marburg University). Jens Sommer, Olaf Steinsträter (Core-Facility Brainimaging, Marburg University). Thomas W.D. Möbius (Institute of Medical Informatics and Statistics, Kiel University).

CP1, biobank. Non-author PIs and members include: Petra Pfefferle (PI; PF 784/1-1, PF 784/1-2), Harald Renz (PI; RE 737/20-1, 737/20-2), Julian Glandorf, Fabian Kormann, Arif Alkan, Fatana Wedi, Lea Henning, Alena Renker, Karina Schneider, Elisabeth Folwarczny, Dana Stenzel, Kai Wenk, Felix Picard, Alexandra Fischer, Sandra Blumenau, Beate Kleb, Doris Finholdt, Elisabeth Kinder, Tamara Wüst, Elvira Przypadlo, Corinna Brehm (Comprehensive Biomaterial Bank Marburg, Marburg University).

## Data access and responsibility

FOR2107/MACS cohort data: All PIs take responsibility for the integrity of the respective study data and their components. All authors and coauthors had full access to all study data.

MNC data: AK and UD take responsibility for the integrity of the data and the accuracy of data analysis.

## Previously published work

Both datasets are part of a larger consortium and have already been published numerous times (please see https://for2107.de/publikationen/ for the MACS and https://www.medizin.uni-muenster.de/translap/studien/nae.html for the MNC). This manuscript presents the first analysis of gray matter volumes of initially healthy individuals with a known future transition to major depressive disorder (converters). We compared these to both, patients with major depressive disorder and sustained healthy controls. To increase the sample size, especially of the converter subsample, we combined datasets of the MACS and the MNC. For the first time, we included only individuals with available gray matter data from the MRI assessment who also completed the 2-year clinical follow-up assessment from both cohorts.

