## Supplements for "Brain Structural Correlates of an Impending Initial Major Depressive Episode"

### Supplemental Information

#### Supplementary Methods

##### *Recruitment and exclusion criteria*

In the MACS cohort, patients with major depressive disorder (MDD) were recruited through local psychiatric hospitals or newspaper advertisements. In the Münster Neuroimaging Cohort (MNC), patients were recruited from the inpatient service of two local psychiatric hospitals. Recruitment of healthy controls (HC) was carried out through public notices and newspaper announcements in both cohorts. Any lifetime diagnoses of schizophrenia, schizoaffective disorder, bipolar disorder, or substance dependence according to DSM-IV (1) were excluded. Further exclusion criteria were neurological abnormalities, a history of seizures, head trauma or unconsciousness, severe physical impairments (e.g., cancer or epilepsy), hypothyroidism without proper medication, claustrophobia, color blindness, and general magnetic resonance imaging (MRI) contradictions (such as ferromagnetic implants or pregnancy).

All participants from the ongoing MNC cohort were re-invited approximately every two years, resulting in multiple available follow-up assessments per person. For the HC and MDD subgroup, baseline data was analyzed. For the converter group, data from the assessment (baseline or a follow-up assessment) before their first depressive episode was selected for analysis.

##### *Computation of medication load index*

Type and dose of psychopharmacological treatment were assessed in structured clinical interviews. First, to measure the total medication load, a strategy described earlier was used (2). Each psychotropic medication was coded as absent=0, low=1 (equal or lower average dose), or high=2 (greater than average dose), relative to the midpoint of the daily dose range recommended by Physician's-Desk-Reference (3). A composite measure of total medication load was calculated for each individual, reflecting dose and variety of different medications taken, by summing all individual medication. Next, the mean of the total medication scores was calculated at both time-points. Finally, this total medication load mean was multiplied with the percentage of days receiving medication during follow-up, to integrate days without the influence of psychopharmacological treatment.

***Detailed description of MRI parameters***

In the Marburg-Muenster Affective Disorders Cohort Study (MACS), T1-weighted high-resolution anatomical data were collected at 3T MRI scanners using three-dimensional (3D) fast gradient echo sequences (MPRAGE) recorded by 3T MRI scanners (Marburg: Tim Trio, Siemens, Erlangen, Germany; Münster: Prisma, Siemens, Erlangen, Germany). Sequence parameters were: 176 sagittal slices, slice gap 0.5mm, TR=1900ms, TE=2.26ms, inversion time=900ms, FA=9°, voxel size=1x1x1mm$^3$ (Marburg) and 192 sagittal slices, slice gap 0.5mm, TR=2130ms, TE=2.28ms, inversion time=900ms, FA=8°, voxel size=1x1x1mm$^3$ (Münster). During data acquisition, the body- coil in Marburg was exchanged resulting in three different scanner settings for all analyses (Marburg body- coil pre, Marburg body- coil post and Münster; 4).

In the MNC, 1-weighted high-resolution anatomical images of the head were acquired (Gyroscan Intera 3T, Philips Medical Systems, the Netherlands) using a three-dimensional fast gradient echo sequence (turbo field echo), repetition time=7.4 ms, echo time=3.4 ms, flip angle=9°, two signal averages, inversion prepulse every 814.5 ms, acquired over a field of view of 256 mm (feet-head) x 204 mm (anterior-posterior) x 160 mm (right-left), frequency encoding in feet to head direction, phase encoding in anterior-posterior and right-left direction, reconstructed to voxels of 0.5 mm × 0.5 mm × 0.5 mm.

***ComBat Harmonization***

To adjust for scanner and site effects, data was harmonized using the MatLab version of ComBat (https://github.com/Jfortin1/ComBatHarmonization/tree/master/Matlab). ComBat, a popular batch correction tool, has been applied in previous neuroimaging studies using different brain structural modalities (5–7). Using an empirical Bayesian framework, ComBat successfully removes variability attributed to scanner and site differences (8,9).

***Matched sample***

To verify whether potential brain structural alterations are present only before the first ever depressive episode or also prior a recurrent episode, we additionally created a smaller sample (*n*=135) matched for age, sex, and scanner variables within each cohort separately (Table S1-S2). Regarding scanner variables, in the MACS cohort we matched for site (Marburg vs. Münster) and body coil change (pre vs. post body coil change). In the MNC, we matched for the year of MRI assessment, since this study is ongoing since 2009 and includes up to seven follow-up assessments per person. Thus this sample included baseline or follow-up data of HC and converters, depending on the time of onset of MDD. Since patients in the MNC were all suffering from an acute moderate or severe depressive episode when initially recruited, the matched sample only included follow-up data of patients who were in remission at time of assessment. Again, data was harmonized using ComBat (https://github.com/Jfortin1/ComBatHarmonization/tree/master/Matlab).

**Table S1.**

*Sample characteristics of the smaller sample matched for sex, age, and scanner variables divided by group*

| Characteristic | CON (*n*=45) *M (SD)* | HC (*n*=45) *M (SD)* | MDD (*n*=45) *M (SD)* | *p*[1] | CON vs. HC *p*[2] | CON vs. MDD *p*[3] |
|---|---|---|---|---|---|---|
| Age | 34.02 (11.46) | 37.31 (13.93) | 35.60 (12.45) | .470 | .225 | .531 |
| Gender (m/f)[4] | 18/27 | 21/24 | 20/25 | .810 | .523 | .670 |
| Cohort (MACS/MNC)[4] | 30/15 | 30/15 | 30/15 | 1.000 | 1.000 | 1.000 |
| HDRS Score | 1.71 (2.50) | 1.09 (2.25) | 6.69 (5.25) | **<.001** | .218 | **<.001** |
| Familial risk for MDD (no risk/risk)[4] | 31/14 | 45/0 | 29/16 | **<.001** | **<.001** | .655 |

*Note.* CON=converters, HC=healthy controls, MDD=major depressive disorder, MACS=Marburg Münster Affective Disorders Cohort Study, MNC=Münster Neuroimaging Cohort, HDRS=Hamilton Depression Rating Scale. Significant p-values ($p < .05$) are highlighted in bold. The matched sample includes remitted patients only who experienced a relapse within a 2-year follow-up interval.

[1] MDD vs. CON vs. HC using one-way ANOVA except where noted.

[2] CON vs. HC using unpaired two-tailed t-test except where noted.

[3] CON vs. MDD using unpaired two-tailed t-test except where noted.

[4] $X^2$-test.

**Table S2.**

*Clinical characteristics of patients with MDD and converters for the smaller sample matched for sex, age, and scanner variables divided by study cohort*

| | Patients with MDD | | | Converters | | |
|---|---|---|---|---|---|---|
| | MACS (*n*=30) | MNC (*n*=15) | | MACS (*n*=30) | MNC (*n*=15) | |
| Characteristic | *M* *(SD)* | *M* *(SD)* | $p^1$ | *M* *(SD)* | *M* *(SD)* | $p^1$ |
| Time since onset (month) | 162.80 (120.69) | 99.66 (58.04) | **.022** | NA | NA | NA |
| Number of depressive episodes lifetime | 4.08 (3.97) | 4.33 (2.69) | .828 | NA | NA | NA |
| Cumulative duration of depression (month) | 74.05 (100.20) | 25.17 (21.45) | **.046** | NA | NA | NA |
| Number of in-patient treatments lifetime | 1.43 (1.59) | 2.00 (1.25) | .235 | NA | NA | NA |
| Cumulative duration of in-patient treatment (weeks) | 10.60 (11.46) | 14.60 (23.67) | .544 | NA | NA | NA |
| Comorbidities (no/yes)[2] | 14/16 | 5/10 | .393 | NA | NA | NA |
| Medication Load Index | 1.20 (1.19) | 1.20 (1.37) | 1.000 | NA | NA | NA |
| Time until onset (month) | NA | NA | NA | 12.29 (7.29) | 8.36 (7.22) | .116 |

*Note.* MACS=Marburg Münster Affective Disorders Cohort Study, MNC=Münster Neuroimaging Cohort MDD=major depressive disorder. Significant p-values ($p < .05$) are highlighted in bold. The matched sample includes remitted patients only who experienced a relapse within a 2-year follow-up interval.

[1] MACS vs. MNC using unpaired two-tailed t-test except where noted.

[2] $X^2$-test

***Sanity checks***

In the converter group, we found a significantly higher ratio of familial risk for MDD compared to HC. Thus, besides controlling for psychopharmacological treatment, we further repeated the ANCOVAs from analysis 1 separately with family risk for MDD as additional covariate to exclude confounding effects (Table S9). The statistical threshold was set to a conservative familywise-error (FWE) correction of $p < 0.05$ on the voxel-level, without cluster extent threshold. To check for influential data points, mean cluster values of the significant amygdala cluster were extracted and checked for outliers using boxplots. The boxplots identified neither extreme outliers with values beyond the cutoff of >3 interquartile ranges across all participants (Figure S2) nor divided by group (Figure S3). Lastly, we checked whether higher gray matter volumes (GMV) in the amygdala in converters were associated with the time until the onset of the initial depressive episode (Figure S4).

#### Supplementary Results

**Table S3.**

*Results from ANCOVAs and t-tests (analyses 1) in the total sample (n=1709) for the dorsolateral prefrontal cortex, insula and amygdala*

| | | | Peak voxel coordinates | | | | | |
|---|---|---|---|---|---|---|---|---|
| Contrast | Side | Cluster size | x | y | z | *F-/t-* value | $p_{TFCE\text{-}FWE}$-value | Effect size partial $\eta^2$/*d* |
| | | | Dorsolateral prefrontal cortex | | | | | |
| Main effect group | R | 92 | 34 | 44 | 30 | 9.37 | **.032** | .011 |
| HC>MDD | R | 825 | 36 | 44 | 32 | 3.81 | **.006** | .188 |
| | R | 37 | 32 | 48 | 16 | 3.41 | **.043** | .168 |
| HC<MDD | - | - | - | - | - | - | .643 | - |
| CON>MDD | - | - | - | - | - | - | .302 | - |
| CON<MDD | - | - | - | - | - | - | .348 | - |
| HC>CON | - | - | - | - | - | - | .157 | - |
| HC<CON | - | - | - | - | - | - | >.999 | - |
| | | | Insula | | | | | |
| Main effect group | R | 14 | 45 | 0 | 2 | 7.40 | **.048** | .009 |
| HC>MDD | R | 703 | 44 | 0 | 2 | 3.78 | **.010** | .186 |
| HC<MDD | - | - | - | - | - | - | .677 | - |
| CON>MDD | - | - | - | - | - | - | .067 | - |
| CON<MDD | - | - | - | - | - | - | .367 | - |
| HC>CON | - | - | - | - | - | - | .170 | - |
| HC<CON | - | - | - | - | - | - | .360 | - |
| | | | Amygdala | | | | | |
| Main effect group | R | 156 | 30 | -3 | -24 | 5.71 | **.007*** | .007 |
| HC>MDD | L | 3 | -21 | 0 | -16 | 2.73 | **.049** | .135 |
| HC<MDD | - | - | - | - | - | - | >.999 | - |
| CON>MDD | R | 283 | 30 | -3 | -24 | 3.31 | **.005** | 0.508 |
| CON<MDD | - | - | - | - | - | - | >.999 | - |
| HC>CON | - | - | - | - | - | - | .195 | - |

| | | | Peak voxel coordinates | | | | | |
|---|---|---|---|---|---|---|---|---|
| Contrast | Side | Cluster size | x | y | z | *F-/t-* value | $p_{TFCE\text{-}FWE}$-value | Effect size partial $\eta^2/d$ |
| HC<CON | R | 64 | 30 | -3 | -24 | 2.93 | **.037** | 0.447 |

*Note.* HC=healthy controls, CON=converters, MDD=major depressive disorder. Significant *p*-values of the significance level $p < .05$ are highlighted in bold.
*Significant after Bonferroni correction for multiple statistical tests (*p*=.01)

**Table S4.**

*Results from ANCOVAs and t-tests (analyses 2) in the remission sample (n=1310) for the dorsolateral prefrontal cortex, insula and amygdala*

| Contrast | Side | Cluster size | x | y | z | *F*-/*t*-value | $p_{TFCE-FWE}$-value | Effect size partial $\eta^2$/*d* |
|---|---|---|---|---|---|---|---|---|
| | | | Peak voxel coordinates | | | | | |
| | Dorsolateral prefrontal cortex | | | | | | | |
| Main effect group | - | - | - | - | - | - | .300 | - |
| HC>MDD | - | - | - | - | - | - | .114 | - |
| HC<MDD | - | - | - | - | - | - | .548 | - |
| CON>MDD | - | - | - | - | - | - | .253 | - |
| CON<MDD | - | - | - | - | - | - | .361 | - |
| HC>CON | - | - | - | - | - | - | .164 | - |
| HC<CON | - | - | - | - | - | - | .905 | - |
| | Insula | | | | | | | |
| Main effect group | - | - | - | - | - | - | .288 | - |
| HC>MDD | - | - | - | - | - | - | .175 | - |
| HC<MDD | - | - | - | - | - | - | .630 | - |
| CON>MDD | - | - | - | - | - | - | .061 | - |
| CON<MDD | - | - | - | - | - | - | .588 | - |
| HC>CON | - | - | - | - | - | - | .159 | - |
| HC<CON | - | - | - | - | - | - | .267 | - |
| | Amygdala | | | | | | | |
| Main effect group | - | - | - | - | - | - | .083 | - |
| HC>MDD | - | - | - | - | - | - | .146 | - |
| HC<MDD | - | - | - | - | - | - | >.999 | - |
| CON>MDD | R | 316 | 28 | -3 | -24 | 3.17 | **.002** | **.012** |
| CON<MDD | - | - | - | - | - | - | .747 | - |
| HC>CON | - | - | - | - | - | - | .194 | - |
| HC<CON | R | 189 | 30 | -3 | -24 | 2.86 | **.014** | **.042** |

*Note.* HC=healthy controls, CON=converters, MDD=major depressive disorder. Significant *p*-values of the significance level $p < .05$ are highlighted in bold.

**Table S5.**

*Results from ANCOVA and t-tests (analyses 3) in the recurrence sample (n=1109) for the amygdala*

| | | | Peak voxel coordinates | | | | | |
|---|---|---|---|---|---|---|---|---|
| Contrast | Side | Cluster size | x | y | z | *F*-/*t*-value | $p_{TFCE\text{-}FWE}$-value | Effect size partial $\eta^2/d$ |
| Main effect group | - | - | - | - | - | - | .151 | - |
| HC>MDD | - | - | - | - | - | - | .684 | - |
| HC<MDD | - | - | - | - | - | - | >.999 | - |
| CON>MDD | R | 23 | 28 | -3 | -24 | 2.67 | .046 | .455 |
| CON<MDD | - | - | - | - | - | - | .589 | - |
| HC>CON | - | - | - | - | - | - | .240 | - |
| HC<CON | R | 146 | 28 | -3 | -24 | 2.83 | .024 | .432 |

*Note.* HC=healthy controls, CON=converters, MDD=major depressive disorder. Significant *p*-values of the significance level $p < .05$ are highlighted in bold.

**Table S6.**

*Results from exploratory whole-brain analyses in the three samples conducted at p < .001, uncorrected, with a cluster threshold of k=200*

| | | | Peak voxel coordinates | | | | |
|---|---|---|---|---|---|---|---|
| Anatomical region | Side | Cluster size[1] | x | y | z | *t-/F-* value | $p_{unc}$-value |
| | | | Total sample ($n$=1709) | | | | |
| *Main effect group* | | | | | | | |
| Middle temporal gyrus, superior temporal gyrus | R | 741 | 56 | -20 | -14 | 12.5 | **<.001** |
| Angular gyrus, middle occipital gyrus | L | 486 | -46 | -70 | 33 | 10.69 | **<.001** |
| | | 271 | -56 | -68 | 15 | 9.21 | **<.001** |
| *HC>MDD* | | | | | | | |
| Inferior temporal gyrus | L | 474 | -52 | -24 | -27 | 4.42 | **<.001** |
| Middle temporal gyrus, superior temporal gyrus, middle temporal pole | R | 1142 | 56 | -21 | -12 | 4.41 | **<.001** |
| Angular gyrus, middle occipital gyrus, inferior parietal gyrus | L | 710 | -46 | -69 | 33 | 4.19 | **<.001** |
| Middle temporal gyrus, superior temporal gyrus | L | 615 | -51 | -24 | -15 | 4.09 | **<.001** |
| Middle temporal gyrus | L | 231 | -57 | -68 | 15 | 3.97 | **<.001** |
| Middle frontal gyrus, superior frontal gyrus | R | 447 | 36 | 44 | 32 | 3.80 | **<.001** |
| Thalamus | L/R | 770 | 4 | -6 | 4 | 3.79 | **<.001** |
| Insula, rolandic operculum | R | 201 | 44 | 0 | 2 | 3.77 | **<.001** |
| Cerebelum | R | 244 | 33 | -74 | -50 | 3.48 | **<.001** |
| *HC<MDD* | - | - | - | - | - | - | n.s. |
| *CON>MDD* | | | | | | | |
| Inferior temporal gyrus, fusiform gyrus | L | 218 | -52 | -51 | -20 | 3.83 | **<.001** |
| Middle cingulate & paracingulate gyri | L/R | 301 | -4 | -21 | 44 | 3.75 | **<.001** |
| Middle temporal gyrus, inferior temporal gyrus, middle | L | 253 | -50 | -68 | 9 | 3.60 | **<.001** |

| | | | Peak voxel coordinates | | | | |
|---|---|---|---|---|---|---|---|
| Anatomical region | Side | Cluster size[1] | x | y | z | *t-/F*-value | $p_{unc}$-value |
| occipital gyrus, inferior occipital gyrus | | | | | | | |
| *CON<MDD* | - | - | - | - | - | - | n.s. |
| *HC>CON* | - | - | - | - | - | - | n.s. |
| *HC<CON* | - | - | - | - | - | - | n.s. |
| | Remission sample[2] (*n*=1310) | | | | | | |
| *Main effect group* | - | - | - | - | - | - | n.s. |
| *HC>MDD* | - | - | - | - | - | - | n.s. |
| *HC<MDD* | - | - | - | - | - | - | n.s. |
| *CON>MDD* | - | - | - | - | - | - | n.s. |
| *CON<MDD* | - | - | - | - | - | - | n.s. |
| *HC>CON* | - | - | - | - | - | - | n.s. |
| *HC<CON* | - | - | - | - | - | - | n.s. |
| | Recurrence sample[3] (*n*=1109) | | | | | | |
| *Main effect group* | - | - | - | - | - | - | n.s. |
| *HC>MDD* | - | - | - | - | - | - | n.s. |
| *HC<MDD* | - | - | - | - | - | - | n.s. |
| *CON>MDD* | - | - | - | - | - | - | n.s. |
| *CON<MDD* | - | - | - | - | - | - | n.s. |
| *HC>CON* | - | - | - | - | - | - | n.s. |
| *HC<CON* | - | - | - | - | - | - | n.s. |

*Note.* HC=healthy controls, CON=converters, MDD=major depressive disorder.
[1] only significant clusters ($p_{unc}$-value < .001) with cluster size $k$≥200 are reported.
[2] Includes remitted MDD patients only.
[3] Includes remitted MDD patients only who experienced a relapse within 2-year follow-up. Matched for sex, age, scanner variables.

**Table S7.**

*Results from ANCOVAs and t-tests in the smaller sample matched for age, sex and scanner variables for the amygdala (n=135)*

| Contrast | Side | Cluster size | x | y | z | *F-/t-* value | $p_{TFCE\text{-}FWE}$-value | Effect size partial $\eta^2/d$ |
|---|---|---|---|---|---|---|---|---|
| | | | Peak voxel coordinates | | | | | |
| Main effect group | R | 60 | 32 | 0 | -27 | 4.10 | **.042** | .060 |
| HC>MDD | - | - | - | - | - | - | .371 | - |
| HC<MDD | - | - | - | - | - | - | .373 | - |
| CON>MDD | R | 226 | 21 | 6 | -18 | 2.78 | **.016** | .490 |
| CON<MDD | - | - | - | - | - | - | >.999 | - |
| HC>CON | - | - | - | - | - | - | >.999 | - |
| HC<CON | - | - | - | - | - | - | .080 | - |

*Note.* HC=healthy controls, CON=converters, MDD=major depressive disorder. Significant *p*-values of the significance level $p < .05$ are highlighted in bold.

**Table S8.**

*Results from exploratory whole-brain analyses in the smaller sample matched for age, sex and scanner variables (n=135) conducted at p<.001, uncorrected, with a cluster threshold of k=200*

| | | | Peak voxel coordinates | | | | |
|---|---|---|---|---|---|---|---|
| Anatomical region | Side | Cluster size[1] | x | y | z | *t-/F-* value | $p_{unc}$-value |
| *Main effect group* | | | | | | | |
| Middle temporal gyrus, middle occipital gyrus | L | 424 | -48 | -68 | 10 | 12.50 | **<.001** |
| Middle temporal gyrus | L | 249 | -60 | -22 | -12 | 3.79 | **<.001** |
| Middle temporal gyrus, superior temporal gyrus | R | 398 | 62 | -6 | -15 | 3.68 | **<.001** |
| *HC>MDD* | | | | | | | |
| Postcentral gyrus, superior parietal gyrus | R | 308 | 40 | -39 | 64 | 4.61 | **<.001** |
| *HC<MDD* | - | - | - | - | - | - | n.s. |
| *CON>MDD* | | | | | | | |
| Middle temporal gyrus, middle occipital gyrus | L | 967 | -48 | -66 | 9 | 4.97 | **<.001** |
| Inferior frontal gyrus, opercular part, inferior frontal gyrus, triangular part | L | 511 | -56 | 20 | 0 | 4.56 | **<.001** |
| Middle temporal gyrus, superior temporal gyrus | R | 1002 | 62 | -6 | -14 | 4.37 | **<.001** |
| Middle temporal gyrus, inferior temporal gyrus | L | 818 | -58 | -27 | -12 | 4.36 | **<.001** |
| Middle temporal gyrus, inferior temporal gyrus | L | 548 | -51 | -48 | -3 | 4.06 | **<.001** |
| Middle cingulate & paracingulate gyri | L | 233 | -6 | -28 | 42 | 3.77 | **<.001** |
| *CON<MDD* | - | - | - | - | - | - | n.s. |
| *HC>CON* | - | - | - | - | - | - | n.s. |
| *HC<CON* | | | | | | | |
| Middle cingulate & paracingulate gyri, precuneus, paracentral lobule | L | 209 | -10 | -36 | 54 | 3.99 | **<.001** |
| Middle temporal gyrus | R | 205 | 64 | -8 | -18 | 3.61 | **<.001** |

*Note.* HC=healthy controls, CON=converters, MDD=major depressive disorder. Includes remitted MDD patients only who experienced a relapse within 2-year follow-up.

**Table S9.**

*Results from additional analyses in the total sample with psychotropic medication and familiar risk for MDD as additional covariates for the dorsolateral prefrontal cortex, insula and amygdala*

| | | | Peak voxel coordinates | | | | | |
|---|---|---|---|---|---|---|---|---|
| Contrast | Side | Cluster size[1] | x | y | z | *t-/F-* value | $p_{TFCE\text{-}FWE}$-value | Effect size partial $\eta^2$/*d* |
| | | Psychiatric medication | | | | | | |
| *DLPFC* | | | | | | | | |
| Main effect group | - | - | - | - | - | - | >.999 | - |
| HC>MDD | - | - | - | - | - | - | .352 | - |
| HC<MDD | - | - | - | - | - | - | .493 | - |
| CON>MDD | R | 346 | 34 | 42 | 30 | 2.42 | **.013** | .371 |
| CON<MDD | - | - | - | - | - | - | .065 | - |
| HC>CON | - | - | - | - | - | - | .194 | - |
| HC<CON | - | - | - | - | - | - | .93 | - |
| *Insula* | | | | | | | | |
| Main effect group | - | - | - | - | - | - | .170 | - |
| HC>MDD | - | - | - | - | - | - | .106 | - |
| HC<MDD | - | - | - | - | - | - | .528 | - |
| CON>MDD | R | 1422 | 46 | 14 | -9 | 3.13 | **<.001** | .480 |
| | L | 1308 | -48 | 3 | 4 | 1.91 | **.004** | .293 |
| CON<MDD | - | - | - | - | - | - | .063 | - |
| HC>CON | - | - | - | - | - | - | .224 | - |
| HC<CON | - | - | - | - | - | - | .272 | - |
| *Amygdala* | | | | | | | | |
| Main effect group | R | 121 | 30 | -3 | -24 | 6.70 | **.027** | .008 |
| HC>MDD | - | - | - | - | - | - | .078 | - |
| HC<MDD | - | - | - | - | - | - | >.999 | - |
| CON>MDD | R | 526 | 30 | -3 | -24 | 3.54 | **<.001** | .543 |
| | L | 276 | -28 | -4 | -24 | 1.71 | **.014** | .262 |

| | | | Peak voxel coordinates | | | | | |
|---|---|---|---|---|---|---|---|---|
| Contrast | Side | Cluster size[1] | x | y | z | *t-/F-* value | $p_{TFCE\text{-}FWE}$-value | Effect size partial $\eta^2$/*d* |
| CON<MDD | - | - | - | - | - | - | .209 | - |
| HC>CON | - | - | - | - | - | - | .264 | - |
| HC<CON | R | 181 | 30 | -3 | -24 | 2.93 | **.017** | .447 |
| | | | Familiar risk for MDD | | | | | |
| *DLPFC* | | | | | | | | |
| Main effect group | R | 122 | 34 | 44 | 30 | 9.57 | **.028** | .011 |
| HC>MDD | R | 1008 | 38 | 44 | 32 | 3.91 | **.004** | .193 |
| HC<MDD | - | - | - | - | - | - | .812 | - |
| CON>MDD | - | - | - | - | - | - | .327 | - |
| CON<MDD | - | - | - | - | - | - | .159 | - |
| HC>CON | - | - | - | - | - | - | .484 | - |
| HC<CON | - | - | - | - | - | - | .397 | - |
| *Insula* | | | | | | | | - |
| Main effect group | R | 52 | 44 | 0 | 3 | 7.22 | **.044** | .008 |
| | R | 8 | 46 | 14 | -9 | 7.11 | **.049** | .008 |
| HC>MDD | R | 937 | 44 | 0 | 2 | 3.75 | **.007** | .185 |
| HC<MDD | - | - | - | - | - | - | .828 | - |
| CON>MDD | - | - | - | - | - | - | .053 | - |
| CON<MDD | - | - | - | - | - | - | .140 | - |
| HC>CON | - | - | - | - | - | - | .591 | - |
| HC<CON | R | 70 | 46 | 15 | -9 | 2.71 | **.026** | .414 |
| *Amygdala* | | | | | | | | |
| Main effect group | R | 68 | 30 | -3 | 24 | 5.73 | **.040** | .007 |
| HC>MDD | L | 47 | -20 | 0 | 16 | 2.87 | **.039** | .141 |
| HC<MDD | - | - | - | - | - | - | >.999 | - |
| CON>MDD | R | 326 | 30 | -3 | -24 | 3.33 | **.002** | .511 |
| CON<MDD | - | - | - | - | - | - | .359 | - |
| HC>CON | - | - | - | - | - | - | .790 | - |

| | | | Peak voxel coordinates | | | | | |
|---|---|---|---|---|---|---|---|---|
| Contrast | Side | Cluster size[1] | x | y | z | *t-/F-* value | $p_{TFCE\text{-}FWE}$-value | Effect size partial $\eta^2$/*d* |
| HC<CON | R | 373 | 30 | -3 | 24 | 2.97 | **<.001** | .453 |

*Note.* HC = healthy controls, CON = converters, MDD = major depressive disorder. Significant *p*-values of the significance level $p < .05$ are highlighted in bold.

**Figure S1**

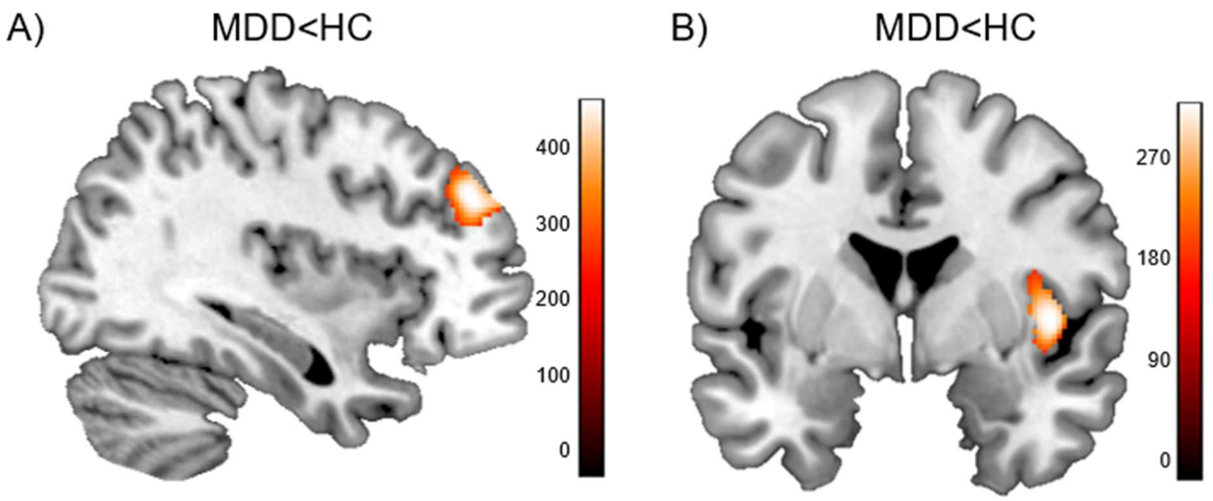


Gray matter volume reductions in the total sample of patients with MDD vs. healthy controls (MDD<HC *t*-contrast) significant at $p_{TFCE\text{-}FWE}$<.05. Depicted are the significant clusters **A)** within the dorsolateral prefrontal cortex ROI at x=222, y=303, z=204 and **B)** within the insula ROI at x=231, y=215, z=144. Color bar: TFCE values.

**Figure S2**

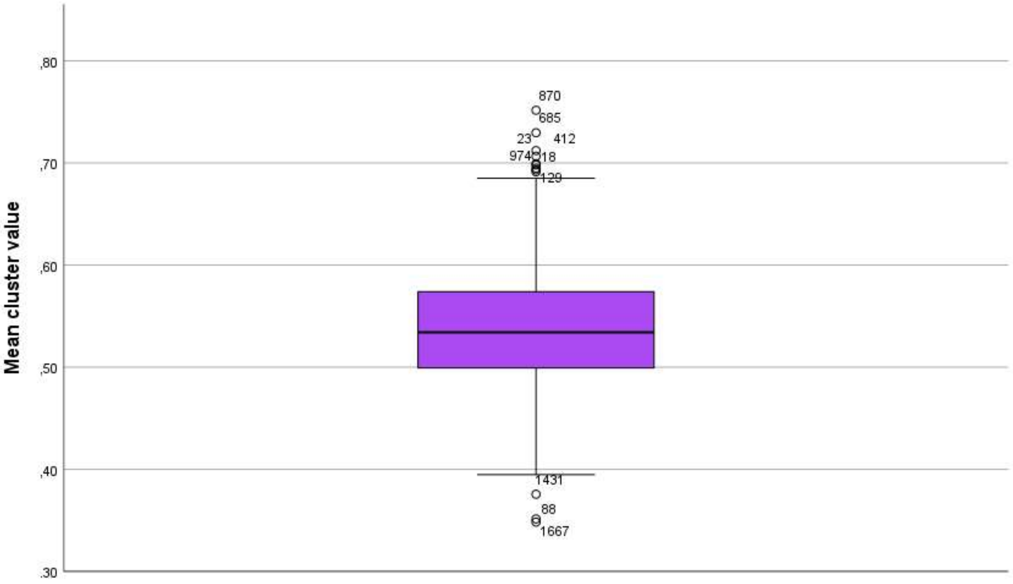


Boxplot of mean cluster values of the significant amygdala cluster (x=30, y=-3, z=-24) across all participants. There were no extreme outliers with values >3 interquartile ranges.

**Figure S3**

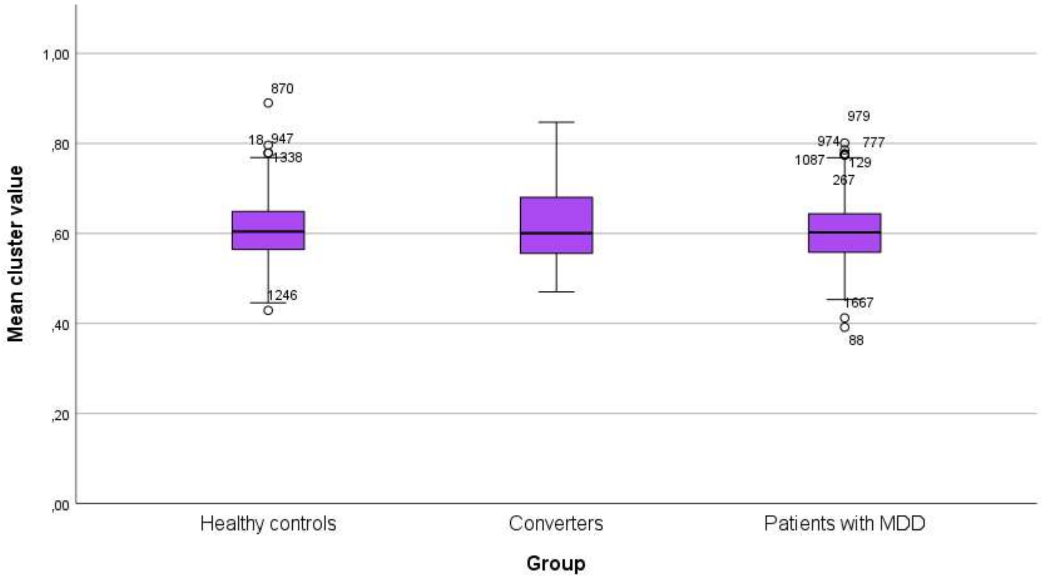


Boxplot of mean cluster values of the significant amygdala cluster (x=30, y=-3, z=-24) divided by group. There were no extreme outliers with values >3 interquartile ranges. MDD=major depressive disorder.

**Figure S4**

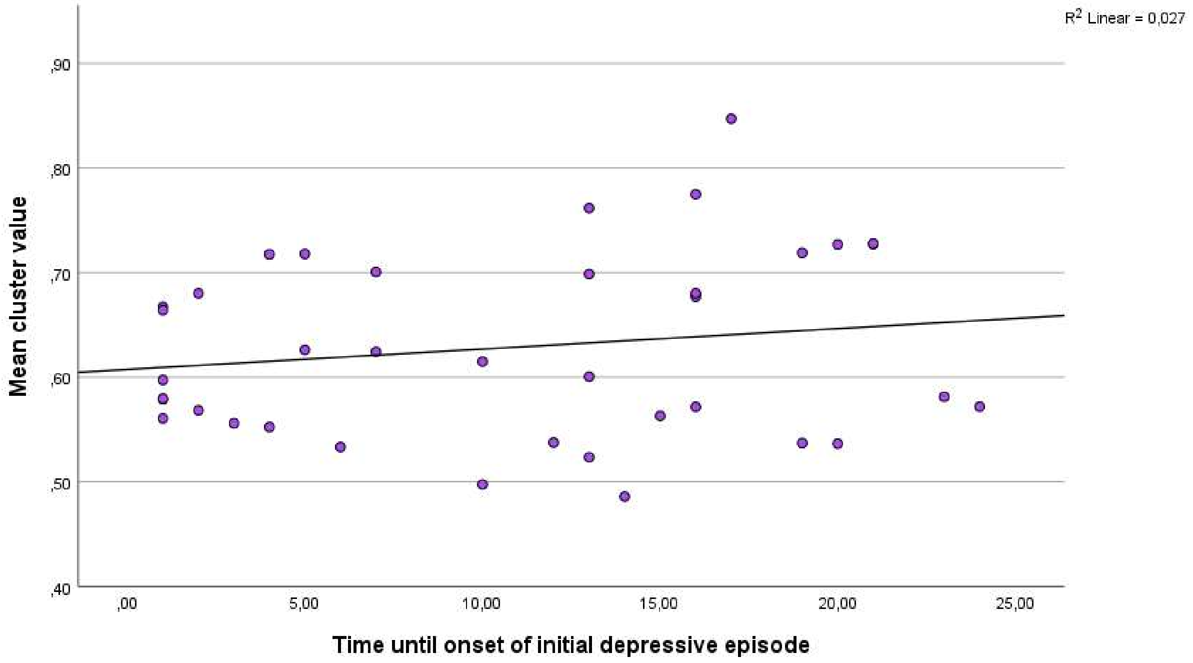


Relation of amygdala gray matter volumes and time until onset of the initial depressive episode in converters (*n*=38). No significant association of mean cluster values of the significant amygdala cluster derived from the F-test in analysis 1 (x=30, y=-3, z=-24) with time until onset of the initial depressive episode in the converter subsample (r=.17, p=.32). For *n*=7 converters information on exact time of onset of the first depressive episode was missing. GMV=gray matter volumes.
